# 6-OHDA-induced dopaminergic neurodegeneration in *C. elegans* is promoted by the engulfment pathway and inhibited by the transthyretin-related protein TTR-33

**DOI:** 10.1101/198606

**Authors:** Sarah-Lena Offenburger, Xue Yan Ho, Theresa Tachie-Menson, Sean Coakley, Massimo A. Hilliard, Anton Gartner

## Abstract

Oxidative stress is linked to many pathological conditions including the loss of dopaminergic neurons in Parkinson’s disease. The vast majority of disease cases appear to be caused by a combination of genetic mutations and environmental factors. We screened for genes protecting *Caenorhabditis elegans* dopaminergic neurons from oxidative stress induced by the neurotoxin 6-hydroxydopamine (6-OHDA) and identified the <u>t</u>rans<u>t</u>hyretin-<u>r</u>elated gene *ttr-33.* The only described *C. elegans* transthyretin-related protein to date, TTR-52, has been shown to mediate corpse engulfment as well as axon repair. We demonstrate that TTR-52 and TTR-33 have distinct roles. TTR-33 is likely produced in the posterior arcade cells in the head of *C. elegans* larvae and is predicted to be a secreted protein. TTR-33 protects *C. elegans* from oxidative stress induced by paraquat or H_2_O_2_ at an organismal level. The increased oxidative stress sensitivity of *ttr-33* mutants is alleviated by mutations affecting the KGB-1 MAPK kinase pathway, whereas it is enhanced by mutation of the JNK-1 MAPK kinase. Finally, we provide genetic evidence that the *C. elegans* cell corpse engulfment pathway is required for the degeneration of dopaminergic neurons after exposure to 6-OHDA. In summary, we describe a new neuroprotective mechanism and demonstrate that TTR-33 normally functions to protect dopaminergic neurons from oxidative stress-induced degeneration, potentially by acting as a secreted sensor or scavenger of oxidative stress.

**Author summary:** Animals employ multiple mechanisms to prevent their cells from damage by reactive oxygen species, chemically reactive molecules containing oxygen. Oxidative stress, caused by the overabundance of reactive oxygen species or a decreased cellular defence against these chemicals, is linked to a variety of neurodegenerative conditions, including the loss of dopaminergic neurons in Parkinson’s disease. In this study, we discovered a novel protective molecule that functions to prevent dopaminergic neurodegeneration caused by oxidative stress induced by the neurotoxin 6-hydroxydopamine (6-OHDA). We used the nematode *C. elegans*, a well-characterised model in which mechanisms can be studied on an organismal level. When *C. elegans* is exposed to 6-OHDA, its dopaminergic neurons gradually die. Our major findings include (i) mutations of the transthyretin-related gene *ttr-33* causes highly increased dopaminergic neurodegeneration after 6-OHDA exposure; (ii) TTR-33 is likely produced and secreted by several cells in the head of the animal; (iii) TTR-33 protects against oxidative stress induced by other compounds; (iv) mutations in the KGB-1 MAP kinase stress pathway alleviate dopaminergic neuron loss in the *ttr-33* mutant; and (v) the cell corpse engulfment pathway is required for dopaminergic neurodegeneration. We hypothesise that TTR-33 protects dopaminergic neurons against 6-OHDA-induced oxidative stress by acting as an oxygen sensor or scavenger.

## Introduction

Oxidative stress, caused by an increased occurrence of reactive oxygen species or a decreased cellular defence against these chemicals, is a risk factor for many pathological conditions including Parkinson’s disease (for review [1]). The aetiology of Parkinson’s disease, which is the second most abundant neurodegenerative disorder, is largely unknown (for review [2]). Although approximately 10% of disease cases can be explained by single gene mutations, the majority are thought to be evoked by the interplay of genetic defects and environmental factors (for review [3,4]). One of the major hallmarks of Parkinson’s disease is the degeneration of dopaminergic neurons in the *substantia nigra* of the midbrain, as well as the abnormal aggregation of intracellular α-synuclein protein known as ‘Lewy bodies’ in surviving dopaminergic neurons [4]. In both genetic and idiopathic cases of Parkinson’s disease, oxidative damage appears to play an important role [4]. Several disease loci are associated with mitochondrial function, and an increased incidence of Parkinson’s is associated with rural living, possibly linked to pesticide exposure [4].

The nematode *C. elegans* is a powerful model system to understand how dopaminergic neurons maintain their integrity when animals are subjected to oxidative stress. Components of dopaminergic neurotransmission are highly conserved between *C. elegans* and mammals, and dopaminergic neurons can be readily visualised in living animals (for review [5]). *C. elegans* hermaphrodites possess eight dopaminergic neurons [6]. These are non-essential for viability or locomotion, whereas they function in mechanosensation and are important for adapting movement to nutrient availability [5].

*C. elegans* dopaminergic neurons have been used to mimic dopaminergic neurodegeneration associated with exposure to known and suspected neurotoxins. One of these toxins is 1-methyl-4- phenyl-1,2,3,6-tetrahydropyridine (MPTP), exposure to which was discovered to cause parkinsonism in a group of drug addicts [7]. In *C. elegans*, MPTP-induced neurodegeneration is independent of the CED-4 apoptosis protein and seems to be in part affected by necrosis [8]. Exposure to manganese also leads to neurodegeneration in *C. elegans* [9,10], as does treatment with paraquat and aflatoxin B1, agents that were recently implicated in dopaminergic neurodegeneration by causing genotoxic damage in mitochondria [11]. One of the most intriguing agents that induces dopaminergic neurodegeneration in *C. elegans* and other model organisms is 6-hydroxydopamine (6-OHDA), a neurotoxin related to dopamine (for review [12]). 6-OHDA is specifically taken up into dopaminergic neurons by the dopamine transporter and leads to oxidative stress by blocking complex I of the respiratory chain [12]. When *C. elegans* are exposed to 6-OHDA, dopaminergic neurons gradually die [13]. It remains underexplored which cell death pathways mediate dopaminergic neurodegeneration after 6-OHDA exposure, and which mechanisms limits their damage upon exposure to oxidative stress.

Key features of apoptotic cell death and cell corpse engulfment mechanisms have been discovered in *C. elegans* (for review [14]). The vast majority of *C. elegans* apoptotic cell deaths are mediated by a conserved pathway comprised of a Apaf-1-like protein CED-4 and the CED-3 caspase (for review [14]). Apoptotic corpses and necrotic cells are engulfed and eliminated by the corpse engulfment pathway [14]. In contrast to mammals, *C. elegans* does not possess specialised phagocytes, but dying cells are engulfed by surrounding cells (for review [15]). In addition, it has been shown that the engulfment pathway cooperates with the apoptotic pathway to ensure efficient apoptosis [16,17]. Furthermore, there are several *C. elegans* cell deaths during which engulfment is used for apoptosis-independent cell killing [18,19]. This type of cell death, recently termed ‘phagoptosis’, also appears to occur in mammalian neurons which are engulfed by surrounding glia cells (reviewed in [20]). When *C. elegans* dopaminergic neurons die in response to 6-OHDA treatment, their neuronal processes exhibit blebbing and are the first to degenerate [13]. Dying cell bodies exhibit condensed chromatin, a common feature of apoptosis; however, blockage of apoptosis does not prevent neuronal death [13]. Typical features of necrotic cell death, such as extensive swelling, do not occur, but dopaminergic neurons were found to be surrounded by glia cells, consistent with a role of phagocytosis in 6-OHDA-induced neurodegeneration [13].

Through an unbiased forward genetic screen, we have discovered that mutations in the *C. elegans* transthyretin-related gene *ttr*-*33* exacerbate dopaminergic neurodegeneration after 6-OHDA treatment. *ttr*-*33* mutants also exhibit increased susceptibility to oxidative stress at an organismal level. *ttr*-*33* is one of the 59 *C. elegans* <u>t</u>rans<u>t</u>hyretin-<u>r</u>elated genes, a family resulting from a nematode-specific expansion [21,22]. Human transthyretin (<u>trans</u>porter of <u>thy</u>roxine (T_4_) and <u>ret</u>inol) is synthesised mainly in the liver and the choroid plexus of the brain, and binds and distributes retinol and thyroid hormones (for review [23]). Transthyretin can form protein aggregates and has been connected to amyloid diseases [23]. The only characterised *C. elegans* transthyretin-related protein, TTR-52, is a secreted protein that facilitates apoptotic corpse engulfment by promoting recognition of apoptotic cells by engulfing cells [24–26]. TTR-52 also mediates axonal repair during regeneration of severed neurons [27]. In contrast, we speculate that TTR-33 may act as a sensor or scavenger of reactive oxygen species. Interestingly, we find that mutations in the *kgb*-*1* MAPK pathway reduce neurodegeneration in *ttr*-*33* mutant animals. Finally, we provide genetic evidence that the *C. elegans* engulfment pathway contributes to the removal of 6-OHDA treated dopaminergic neurons.

## Results

### Mutations in the *C. elegans* transthyretin-related gene *ttr*-*33* causes enhanced 6-OHDA-induced dopaminergic neurodegeneration

Mutations in neuroprotective genes are expected to cause increased neuronal loss after oxidative stress exposure. To discover genes that prevent dopaminergic neurodegeneration, we used a *C. elegans* strain carrying the *vtIs1* transgene, in which the promoter of the dopamine transporter *dat*-*1* drives GFP expression and thus allows selectively visualising the dopaminergic neurons (Fig 1A). These animals (P0) were mutagenised with ethyl methanesulfonate, and L1 stage larvae of the F2 generation were exposed to the neurotoxin 6-hydroxydopamine (6-OHDA) for 1 hour at 10 mM, a treatment that does not induce dopaminergic neurodegeneration in wild-type animals. After 72 hours, the adult F2 population was screened for animals that exhibited excessive loss of dopaminergic head neurons (Fig 1A). We isolated the *gt1983* mutant, which exhibited highly increased 6-OHDA-induced neurodegeneration compared to wild-type animals (Fig 1B). We found that the *gt1983* allele is recessive as *gt1983/wild-type(+)* heterozygous mutants did not exhibit increased 6-OHDA-induced neurodegeneration (Fig 1D). To identify the corresponding locus to the *gt1983* mutation, we performed whole-genome sequencing and single nucleotide polymorphism mapping [28,29]. We found that the *gt1983* mutant carries a point mutation in *ttr*-*33* (*<u>t</u>rans<u>t</u>hyretin-<u>r</u>elated family domain*) that causes a glycine to glutamate conversion (G27E) in the TTR-33 protein (Fig 1C). Another *ttr*-*33* point mutation, *gk567379*, which induces a leucine to phenylalanine substitution (L72F) (Fig 1C), also caused hypersensitivity to 6-OHDA, albeit to a weaker extent (Fig 1E). Furthermore, *gt1983/ttr-33*(*gk567379*) trans-heterozygous animals still exhibited increased 6- OHDA-induced neurodegeneration (Fig 1E). Finally, transgenic expression of wild-type *ttr*-*33* under the control of a 1.5 kb endogenous regulatory region was able to partially rescue the 6-OHDA-induced neurodegeneration defect of *ttr*-*33* mutants (Fig 1F). Taken together, our results indicate that loss-of-function mutations in the transthyretin-related gene *ttr*-*33* lead to increased 6-OHDA-induced dopaminergic neurodegeneration.

**Fig. 1.**
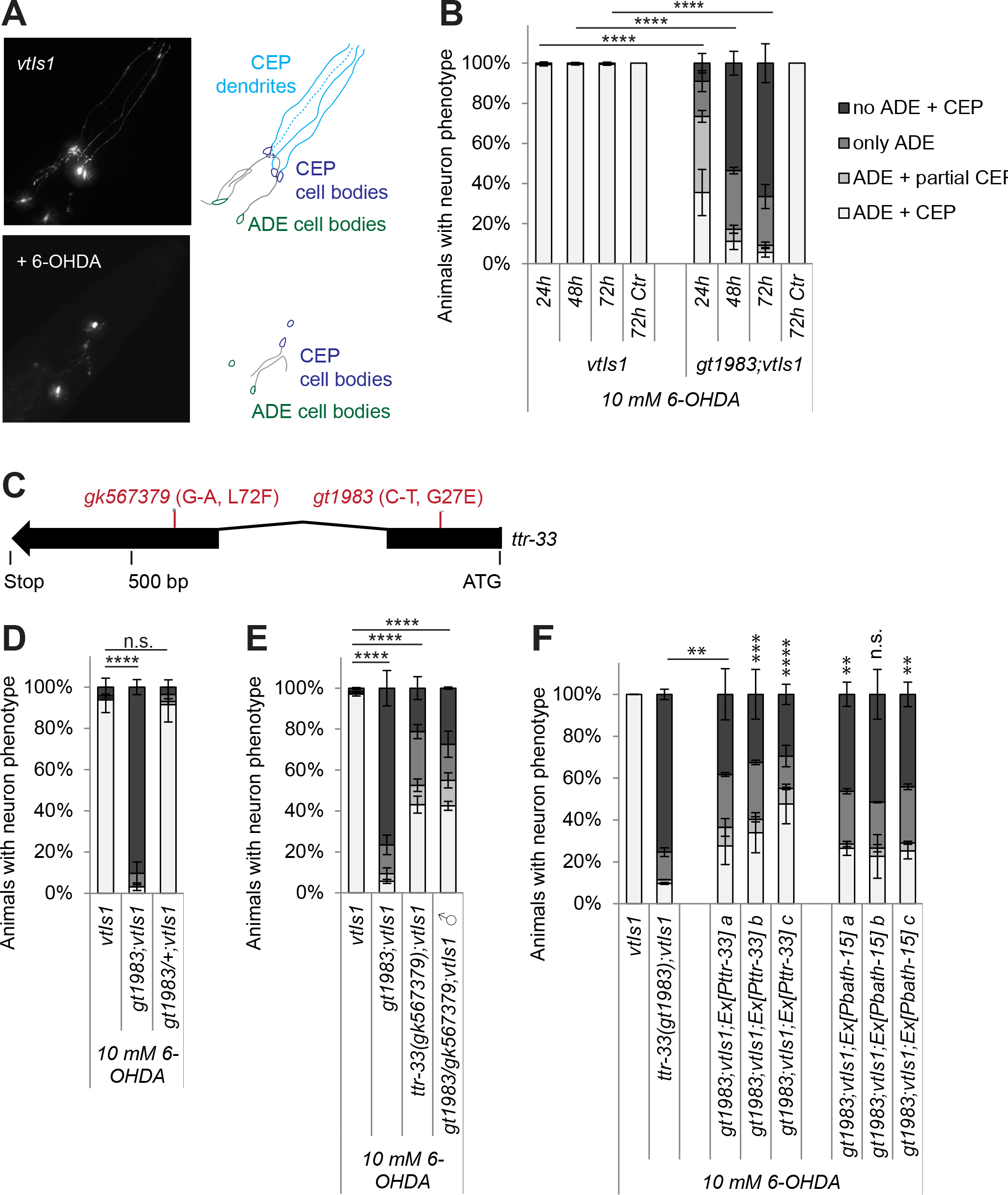
*ttr*-*33* mutations cause increased dopaminergic neurodegeneration after treatment with 6-OHDA. (A) GFP-labelled *C. elegans* dopaminergic head neurons – 4 <u>cep</u>halic sensilla (CEP) neurons and 2 <u>a</u>nterior <u>de</u>irid (ADE) neurons – in animals carrying the *vtIs1* transgene and neurodegeneration after addition of 6-OHDA. (B) Remaining dopaminergic head neurons 24, 48 and 72 hours after treatment with 10 mM 6-OHDA and 72 hours after control treatment with ascorbic acid only (’72h Ctr‘) for BY200 wild-type and *gt1983* mutant animals. Animals possessing all neurons were scored as ‘ADE + CEP’ (white bar), those with partial loss of CEP but intact ADE neurons as ‘ADE + partial CEP’ (light grey bar), those with complete loss of CEP but intact ADE neurons as ‘only ADE’ (dark grey bar) and those with complete loss of dopaminergic head neurons as ‘no ADE + CEP’ (black bar). Error bars; = S.E.M. of 3 biological replicates for all the other strains, each with 60-120 animals per strain. Total number of animals per condition n = 30 for the ‘72h Ctr’ and n = 220-340 for all the other conditions (****p<0.0001; G-Test comparing BY200 wild-type and mutant data of the same time point). (C) *ttr*-*33* gene structure with positions of *ttr*-*33(gt1983)* and *ttr*-*33(gk567379)* point mutations. Nucleotide and amino acid changes are indicated in parentheses. (D) Dopaminergic head neurons after treatment with 10 mM 6-OHDA in BY200 wild-type animals and *ttr*-*33(gt1983)* homozygous and *ttr*-*33*(*gt1983*)/+ heterozygous mutants. Error bars = SEM (Standard Error of the Mean) of 2 experiments, each with 50-105 animals per strain. Total number of animals per strain n = 120-205 (****p<0.0001, n.s. p>0.05; G-Test). (E) Dopaminergic head neurons after treatment with 10 mM 6-OHDA in BY200 wild-type animals, *gt1983* and *ttr*-*33(gk567379)* homozygous mutants and *gt1983*/*ttr*-*33(gk567379)* trans heterozygous mutant animals. Error bars = SEM of 2-3 experiments, each with 100-175 animals per strain. Total number of animals per strain n = 210-340 (n.s. p>0.05; G-Test). (F) Effect of *Pttr-33::ttr*-*33* and *Pbath*-*15::ttr*-*33* extrachromosomal arrays on dopaminergic neurodegeneration of *ttr*-*33* mutant animals after treatment with 10 mM 6-OHDA. Error bars = SEM of 2 biological replicates, each with 90-130 animals per strain and concentration. Total number of animals per condition n = 200-245 (***p<0.001, **p<0.01, *p<0.055, n.s. p>0.05; G-Test). At present, we speculate that the *ttr*-*33* mutant defect might not be fully rescued by our *ttr*-*33* wild-type plasmids because the constructs might not contain all promoter elements. Also, a 3 amino acid linker sequence, introduced as part of the cloning strategy between the translational start site and the N-terminal secretion signal of TTR-33, may compromise protein function.

### 6-OHDA sensitivity of *ttr*-*33* mutant hypersensitivity occurs in the first larval stages and is inhibited by blocking neuronal drug uptake

TTR-33, like the majority of the *C. elegans* transthyretin-related proteins [24], is predicted to be a secreted, non-cytoplasmic protein (S2A Fig). Apart from the small alpha-helical signal peptide at the N-terminus, the predicted TTR-33 protein structure is composed predominantly of beta sheets (Fig 2A; S2C Fig). According to this protein structure prediction, *ttr-33(gt1983)* and *ttr*-*33(gk567379)* lead to amino acid changes on two neighbouring beta-strands (Fig 2A).

**Fig. 2.**
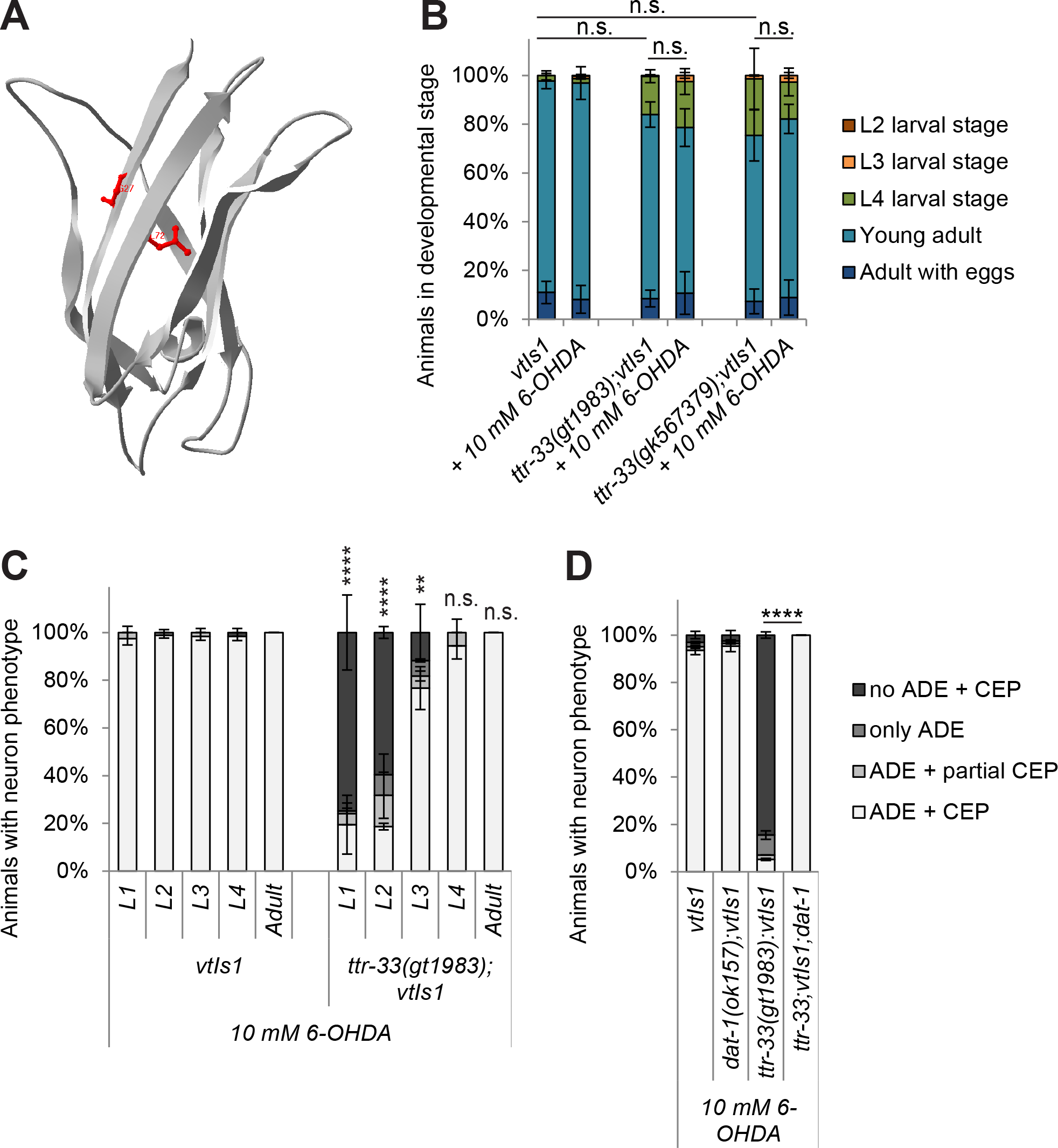
*ttr*-*33* mutants develop at a normal speed and 6-OHDA must enter dopaminergic neurons to cause dopaminergic neurodegeneration. (A) TTR-33 protein structure prediction based on homology modelling using the structure of TTR-52 (PDB ID: 3UAF). The overlay of the TTR-33 and the TTR-52 structures is shown in S1 Fig B. The *ttr*-*33(gt1983)* point mutation leading to a glycine to glutamate (G27E) conversion and the *ttr*-*33(gk567379)* mutation leading to a leucine to phenylalanine (L72F) conversion are both indicated in red. (B) Developmental stages of wild-type and *ttr*-*33* mutant L1 stage larvae 48 hours after treatment with and without 10 mM 6-OHDA. *C. elegans* L1 stage larvae develop via the L2 (in red), L3 (in orange), L4 (green) and young adult stage (in light blue) into adults (in dark blue). Error bars = SEM of 3 biological replicates, each with 60-175 animals per treatment and strain. Total number of animals per condition n = 205-350 (n.s. p>0.05; G-Test). (C) Dopaminergic head neurons in wild-type animals and *ttr*-*33* mutants 48 hours after treatment of L1-L4 larval stages or adult animals with 10 mM 6-OHDA. Error bars = SEM of 2 biological replicates, each with 25-45 animals per stage and strain. Total number of animals per condition n = 50-80 (****p>0.0001, **p<0.01, n.s. p>0.05; G-Test comparing BY200 wild-type and mutant animals data of the same lifecycle stages). (D) Effect of *dat*-*1* mutation on dopaminergic neurodegeneration after treatment with 10 mM 6-OHDA. Error bars = SEM of 2 experiments, each with 90-115 animals per strain. Total number of animals per strain n = 200-535 (****p<0.0001; G-Test).

To determine the developmental stages in which *ttr*-*33* functions to protect *C. elegans* from 6-OHDA-induced neurodegeneration, we exposed a mixed population of *ttr*-*33* mutants to this drug. The larval stages 1-4 (L1-L4) and adult animals were separated after treatment. We found that L1 and L2 *ttr*-*33* mutant larvae were highly sensitive to 6-OHDA, whereas L3 larvae exhibited less sensitivity (Fig 2C). Mutant L4 larvae and adults did not show increased neurodegeneration (Fig 2C). We conclude that loss of function of *ttr*-*33* causes 6-OHDA-induced dopaminergic neurodegeneration in the first three larval stages of *C. elegans*.

*ttr*-*33* mutant animals develop at a slightly slower rate than wild-type animals (S3A Fig), but do not exhibit increased growth delays after exposure to 6-OHDA (Fig 2B). Thus, *ttr*-*33* mutants exhibit highly increased dopaminergic neurodegeneration at a concentration of 6-OHDA that does not affect their overall development. We also examined whether the *ttr*-*33* mutation affects dopaminergic signalling, yet we found that *ttr*-*33(gt1983)* mutants behave like wild-type animals in assays testing dopamine-mediated behaviours (S3A and S3B Fig). Finally, *ttr*-*33* mutants do not exhibit loss of dopaminergic neuron when the uptake of 6-OHDA into these cells is prevented by deletion of the <u>d</u>op<u>a</u>mine <u>t</u>ransporter *dat-1* (Fig 2D).

### TTR-33 acts independently of TTR-52

We next asked whether TTR-33 and the previously characterised TTR-52 [24–26] have analogous roles in neuroprotection and axon repair. To test genetic interactions, we also performed experiments with lower concentrations of 6-OHDA to be able to detect possible enhanced neurodegeneration due to synergic effect (S4A Fig). We found that a *ttr*-*52* loss-of-function mutation did not cause increased 6-OHDA-induced neurodegeneration and did not influence the sensitivity of *ttr*-*33* mutants or wild-type animals (Fig 3A, B; S4B and S4C Fig). Thus, TTR-52 does not appear to play a role in 6-OHDA-induced dopaminergic cell loss. In addition, we performed laser axotomy on the PLM (<u>p</u>osterior <u>l</u>ateral <u>m</u>icrotubule cells) mechanosensory neurons of wild-type and *ttr*-*33* mutant animals, using animals carrying the *zdls5* transgene that express GFP under the control of the mechanosensory neuron-specific *mec*-*4* promoter. We found that PLM axons of *ttr*-*33* mutants did not show overt defects in the rate of axonal regrowth, reconnection or fusion after injury (Fig 3C and C′), unlike *ttr*-*52* mutants which have been shown to present a defect in axonal fusion [27]. In summary, TTR-52 and TTR-33 function appear to have specific and independent functions.

**Fig. 3.**
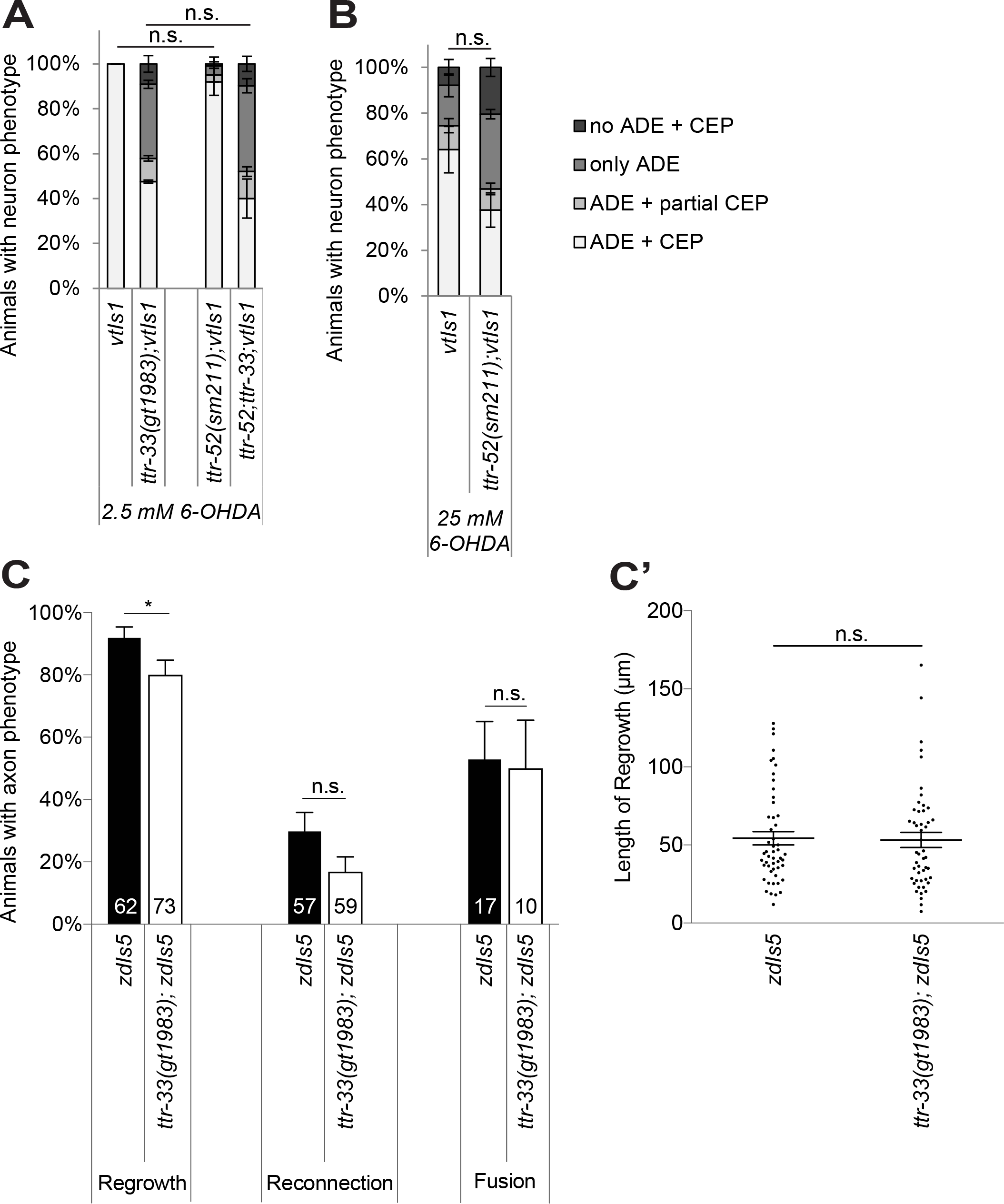
*ttr*-*33* and *ttr*-*52* have independent functions. (A) Effect of *ttr*-*52* mutation on dopaminergic neurodegeneration after treatment with 2.5 mM 6-OHDA. Error bars = SEM of 2 biological replicates, each with 95-115 animals per strain. Total number of animals per condition n = 200-225 (n.s. p>0.05; G-Test). (B) Effect of *ttr*-*52* mutation on dopaminergic neurodegeneration after treatment with 25 mM 6-OHDA. Error bars = SEM of 4 biological replicates, each with 100-130 animals per strain. Total number of animals per condition n = 425-460 (n.s. p>0.05; G-Test). (C) Quantification of axonal regrowth, regeneration and fusion of the PLM axons 24 hours after UV-laser axotomy in *ttr*-*33(gt1983)* mutants and wild-type animals carrying the *zdIs5* transgene to visualise PLM. The percentage of successful reconnection is a proportion of the axons that showed regrowth. The percentage of successful axonal fusion is a proportion of the axons that successfully reconnected. Error bars = standard error of proportion. The number of animals tested is indicated at the bottom of each bar (*p<0.05, n.s. p>0.05; t-test). (C’) Length of axonal regrowth in *ttr-33(gt1983)* mutants and wild-type animals (n.s. p>0.05; t-test).

### The cell corpse engulfment pathway is required for 6-OHDA-induced dopaminergic neuron death

Given the reported role of TTR-52 in corpse engulfment, we asked if the cell corpse engulfment machinery influences 6-OHDA-induced neurodegeneration. The phagocytosis pathway consists of two partially redundant branches (for a simplified model see Fig 4A which was adapted from [27]). One branch involves the TTR-52 bridging molecule, the CED-1 receptor and the CED-6 adaptor protein. The second pathway involves the CED-2 SH2 and SH3 domain-containing adaptor protein, and the activation of the CED-10/RAC protein via the bipartite CED-5 and CED-12 guanine nucleotide exchange factor. CED-10 acts as the end point of both pathways (Fig 4A). We found that mutation in *ced*-*6*, in contrast to mutation in *ced*-*2*, led to a measurable suppression of dopaminergic neurodegeneration in the *ttr*-*33* mutant background (Fig 4B; S5A Fig). Furthermore, *ced*-*2*;*ced*-*6*;*ttr*-*33* triple mutants and *ced*-*10*;*ttr*-*33* double mutants in which both pathways are disrupted, showed an increased suppression of dopaminergic neuron death compared with *ced*-*6*;*ttr*-*33* alone (Fig 4B). Finally, we also observed a reduction in dopaminergic neuron loss in a wild-type background (Fig 4C; S5B Fig). Thus, both branches of the cell corpse engulfment pathway are required for 6-OHDA-induced dopaminergic neuron loss. We note that TTR-52, which has a role in engulfing apoptotic cells in the CED-5 pathway, had no discernible role in 6-OHDA-induced dopaminergic neurodegeneration.

**Fig. 4.**
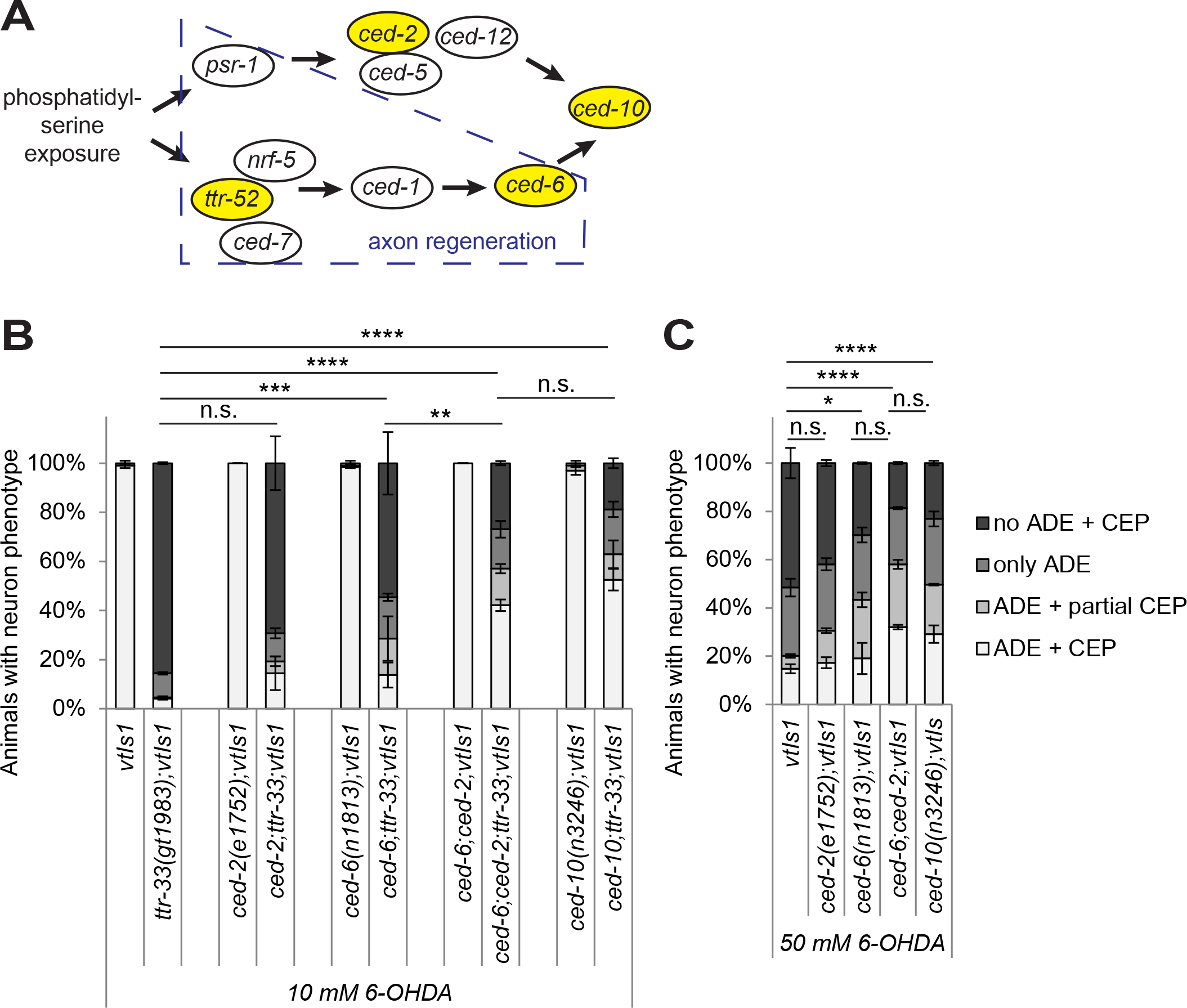
Mutations in the cell corpse engulfment pathway alleviate 6-OHDA-induced dopaminergic neurodegeneration. (A) Cartoon of the engulfment pathway (adapted from [27]). The recognition of the conserved ‘eat-me’ signal phosphatidylserine (PS) on the surface of apoptotic and necrotic cells triggers two partially redundant pathways in the engulfing cell [73]. The genes tested in this study are highlighted in yellow and the part of the pathway that was shown to be required for axon regeneration [27] is boxed in blue. *psr = phosphatidylserine receptor family, ced = cell death abnormality, nrf = nose resistant to fluoxetine (lipid-binding protein).* (B) Effect of mutations in engulfment pathway genes on dopaminergic neurodegeneration after treatment with 10 mM 6-OHDA. Error bars = SEM of 2 biological replicates, each with 70-120 animals per strain. Total number of animals per condition n = 175-230 (****p<0.0001, ***p<0.002, **p<0.01, n.s. p>0.05; G-Test comparing *ttr*-*33* mutant data to double mutant data). (C) Effect of mutations in engulfment pathway genes on dopaminergic neurodegeneration after treatment with 50 mM 6-OHDA. Error bars = SEM of 2 biological replicates, each with 85-140 animals per strain. Total number of animals per condition n = 225-235 (****p<0.0001, *p<0.05, n.s. p>0.05; G-Test).

### *ttr*-*33* is likely expressed in pharynx, hypodermis and in posterior arcade cells

TTR-33 is predicted to be a secreted protein and a transcriptional reporter is thus expected to highlight the location of protein production. We examined three independent transgenic strains carrying transcriptional *Pttr*-*33::gfp* reporter, and found that six cells residing in the head of L1 larvae were strongly labelled (Fig 5A–B). Two of the labelled cells were located next to the first pharyngeal bulb and four were situated halfway between the mouth and the first pharyngeal bulb. All six cells form protrusions towards the mouth of the larva. The cells were labelled as early as in the embryonic pretzel stage (Fig 5C; S1 Movie). Comparison with the cell atlas of *C. elegans* [30] and a GFP reporter under the control of the *grl*-*27* arcade cell-specific promoter [31] revealed that these cells represent most likely the posterior arcade cells, which are located between external epidermis and anterior pharynx [32]. In support of this notion, we found that expression of wild-type TTR-33 under the arcade cell-specific promoter of *bath*-*15 (C08C3.2)* led to a partial rescue of the defect of *ttr*-*33* mutant animals (Fig 1F). In addition, in L1 stage larvae, *ttr*-*33* expression was observed in the pharynx and in several unidentified cells immediately posterior to the second pharyngeal bulb. In older animals, the *Pttr*-*33*::*GFP* transcriptional reporter highlights head cells next to and anterior of the first pharyngeal bulb, as well as the pharynx and the hypodermis (S2-S5 Movie). Given that expression of wild-type TTR-33 under the control of its endogenous regulatory region as well as the *Pbath*-*15* promoter was not sufficient to fully rescue the enhanced neurodegeneration in *ttr*-*33* mutants, it remains possible that the endogenous gene is expressed in further tissue not visible in these transgenic lines.

**Fig. 5.**
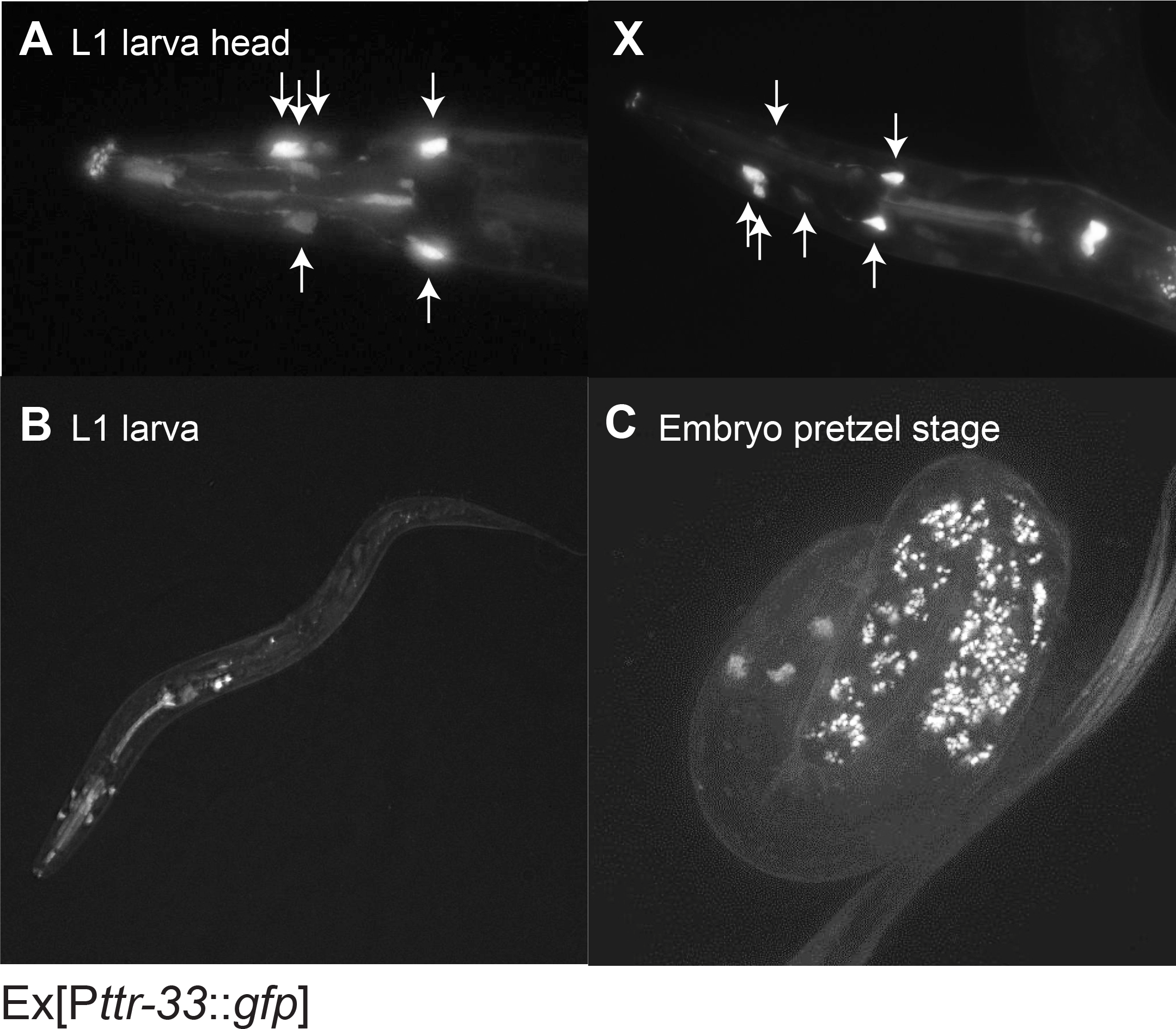
Evidence that *ttr*-*33* is expressed in the posterior arcade cells and in the pharynx. (A) Head region of L1 stage larva expressing the extrachromosomal transcriptional reporter Ex[P*ttr*-*33*::*gfp*]. Arrows indicate strongly expressing cells. (A’) Different focal planes of the same larva. (B) L1 stage larva expressing the transcriptional reporter Ex[P*ttr*-*33*::*gfp*]. (C) Pretzel stage embryo expression the transcriptional reporter Ex[P*ttr*-*33*::*gfp*]. The speckled signal is caused by intestinal autofluorescence.

### *ttr*-*33* mutants are sensitive to oxidative stress

To determine if *ttr*-*33* mutants are also sensitive to oxidative stress at an organismal level, we exposed the L1 stage larvae to different concentrations of paraquat and hydrogen peroxide (H_2_O_2_) for 1 hour. We found that after treatment with paraquat (Fig 6A) and H_2_O_2_ (Fig 6B), the development of *ttr*-*33* mutants is compromised compared with wild-type animals. Since increased oxidative stress has been connected to premature aging (for review [1,33]), we asked whether the lifespan of *ttr*-*33* mutants was altered and found that these animals are indeed short-lived (Fig 6C, S6 Fig). We next investigated if *ttr*-*33* is transcriptionally regulated in response to oxidative stress using qRT-PCR and found that this does not appear to be the case (S7A, I Fig; S8A, I Fig). Furthermore, to determine if oxidative stress signalling is altered in *ttr*-*33* mutants as compared to wild-type animals, we measured expression of the oxidative stress reporters *gst*-*1*, *gst*-*4* and *gcs*-*1*. We found that the *ttr*-*33(gt1983)* mutant does not show overt differences in *gst*-*1*, *gst*-*4* and *gcn-1* mRNA expression after treatment with 6-OHDA and paraquat or under control conditions (S7 Fig and S8 Fig). In summary, TTR-33 appears to provide a constitutive organism-wide oxidative stress protection.

**Fig. 6.**
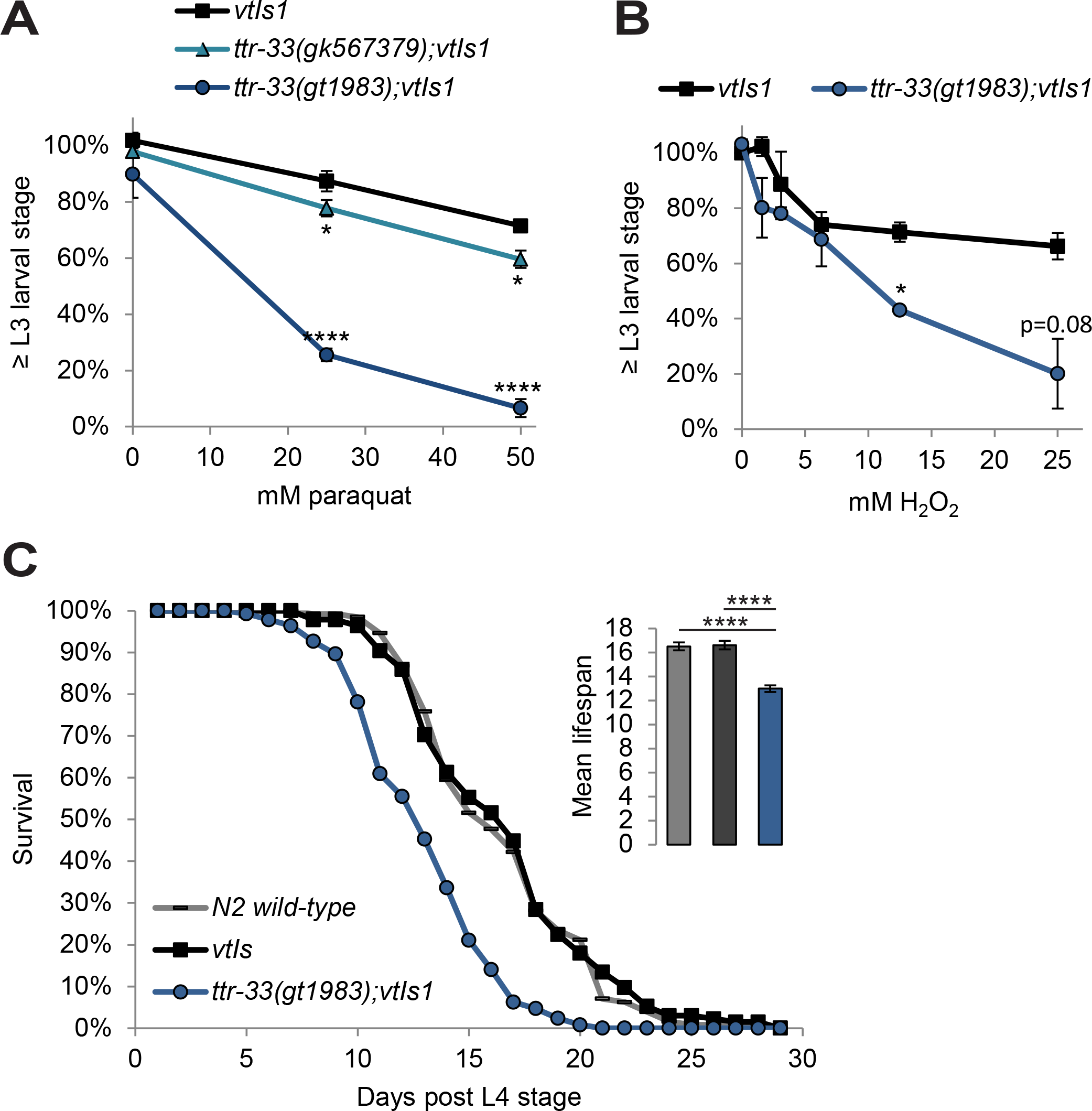
*ttr*-*33* mutants are sensitive to oxidative stress. (A) Percentage of larvae reaching the L3 larval stage 24 hours after an 1 hour incubation with indicated concentration of paraquat. Error bars = SEM of 3 biological replicates for 25 and 50 mM paraquat and 2 biological replicates for 0 mM paraquat, each with 60-245 animals per strain and concentration. Total number of animals per condition n = 275-550 (****p<0.0001, *p<0.05; twotailed t-test comparing BY200 wild-type and mutant animal data). (B) Percentage of animals developed to L3 stage 48 hours after an 1 hour incubation with indicated concentration of H_2_O_2_. Error bars = SEM of 2 biological replicates, each with 165-547 animals per strain and concentration. Total number of animals per condition n = 311-942 (*p<0.05; two-tailed t-test comparing data of BY200 wild-type and mutant animal data at 12.5 and 25 mM H2O2). (C) Lifespan data for first biological replicate including 110-125 animals per strain. The inset shows the mean lifespan with the error bars depicting the standard error. Total number of animals for N2 wild-type (which was tested once) n = 110, and for all other strains n = 200-235 (****Bonferroni p<0.0001; Log-Rank Test).

### JNK stress kinases have a role in 6-OHDA-induced neurodegeneration in *ttr*-*33* mutants

Various MAP kinase pathways are involved in stress signalling (for review [34]), and we therefore tested their role in mediating the response to 6-OHDA. We found that mutations in the p38 kinases PMK-1 (<u>p</u>38 <u>M</u>AP <u>k</u>inase family) and PMK-3 alone did not affect the extent of neurodegeneration (Fig 7). However, a possible redundant function of these two proteins cannot be excluded. In contrast, we found that a *jnk*-*1* (<u>J</u>un <u>N</u>-terminal <u>k</u>inase) mutation led to increased dopaminergic neuron loss in the *ttr*-*33* mutant (Fig 7A; S9A Fig), but not in wild-type animals (Fig 7B, S9B Fig). Interestingly, mutations in the JNK kinase KGB-1 (*<u>k</u>inase*, *<u>G</u>LH*-<u>b</u>inding) strongly alleviated 6-OHDA-induced dopaminergic neurodegeneration in *ttr*-*33(gt1983)* mutants (Fig 8A; S10A Fig) but not in wild-type animals (Fig 8B, S10B Fig). Mutation of MEK-1, the mitogen-activated protein kinase kinase (MAPKK) acting upstream of KGB-1 [34], equally reduced 6-OHDA-induced dopaminergic neuron loss in the *ttr*-*33* mutant (Fig 8A; S10A Fig), but not in wild-type animals (Fig 8B, S10B Fig). Furthermore, we found that mutations in *kgb*-*1* and *mek*-*1* decreased paraquat-induced developmental delays in *ttr*-*33* mutants (Fig 8C). However, *kgb*-*1* and *mek-1* mutations alone did not induce paraquat sensitivity (Fig 8C). Although *mek*-*1* mutation protects from oxidative stress sensitivity in the *ttr*-*33* mutants, *mek*-*1*;*ttr*-*33* double mutants develop slowly and show reduced proliferation (S10 Fig C, D). In summary, the KGB-1 MAP kinase pathway appears to mediate the hypersensitivity to oxidative stress associated with *ttr*-*33* mutation

**Fig. 7.**
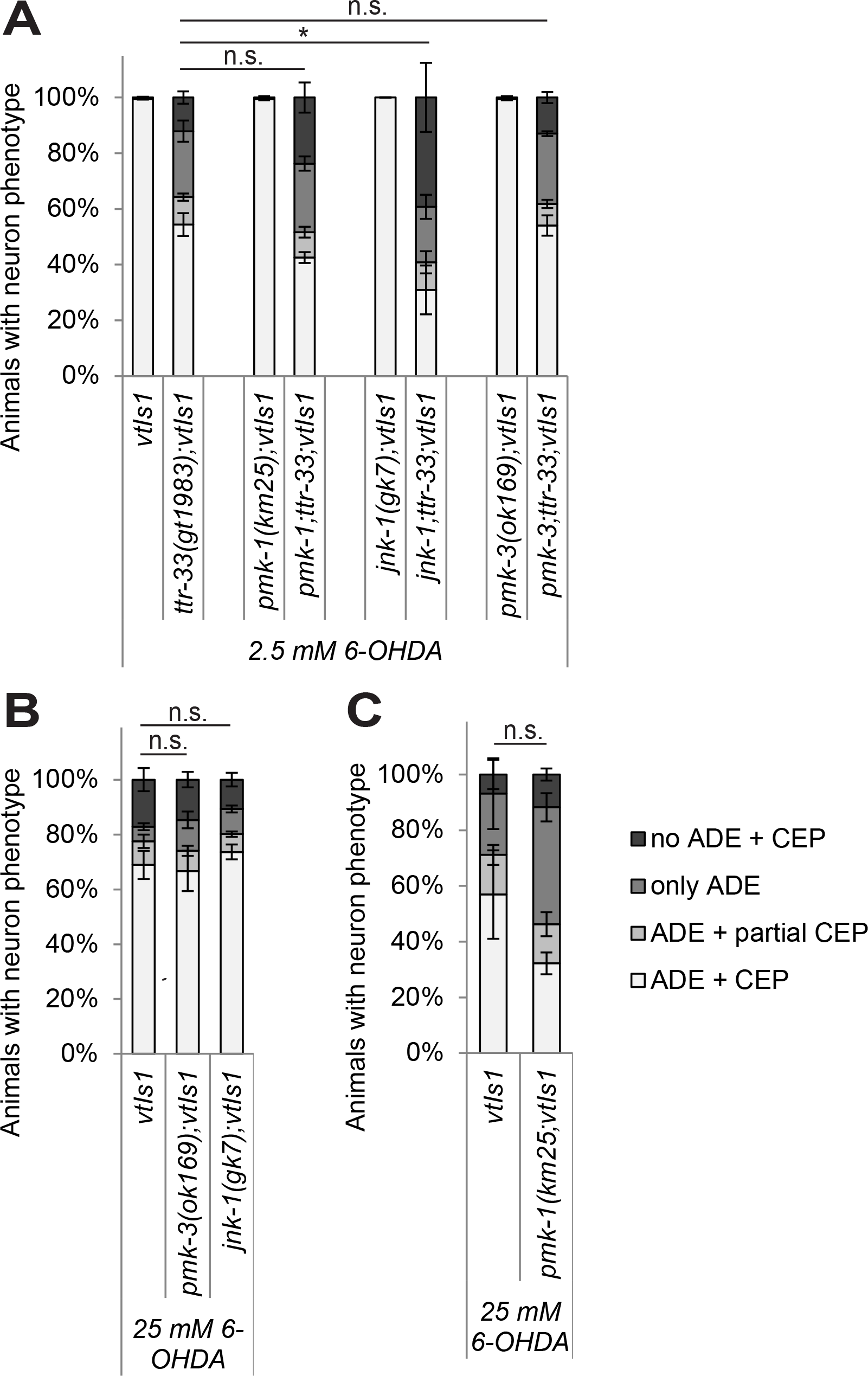
Interference with the *jnk*-*1* MAPK pathway increases 6-OHDA sensitivity in *ttr*-*33* mutants. (A) Effect of p38 and JNK stress response pathway mutations on dopaminergic neurodegeneration after treatment with 2.5 mM 6-OHDA. Error bars = SEM of 2-4 biological replicates, each with 100120 animals per strain and concentration. Total number of animals per condition n = 200-425 (*p<0.05, n.s. p>0.05; G-Test). (B) Effect of p38 and JNK stress response pathway mutations on dopaminergic neurodegeneration after treatment with 25 mM 6-OHDA. Error bars = SEM of 3 biological replicates, each with 100-110 animals per strain. Total number of animals per strain n = 310-330 (n.s. p>0.05; G-Test). (C) Effect of *pmk*-*1* mutation on dopaminergic neurodegeneration after treatment with 25 mM 6-OHDA. Error bars = SEM of 3 biological replicates, each with 100-135 animals per strain. Total number of animals per strain n = 350-360 (n.s. p<0.05; G-Test).

**Fig. 8.**
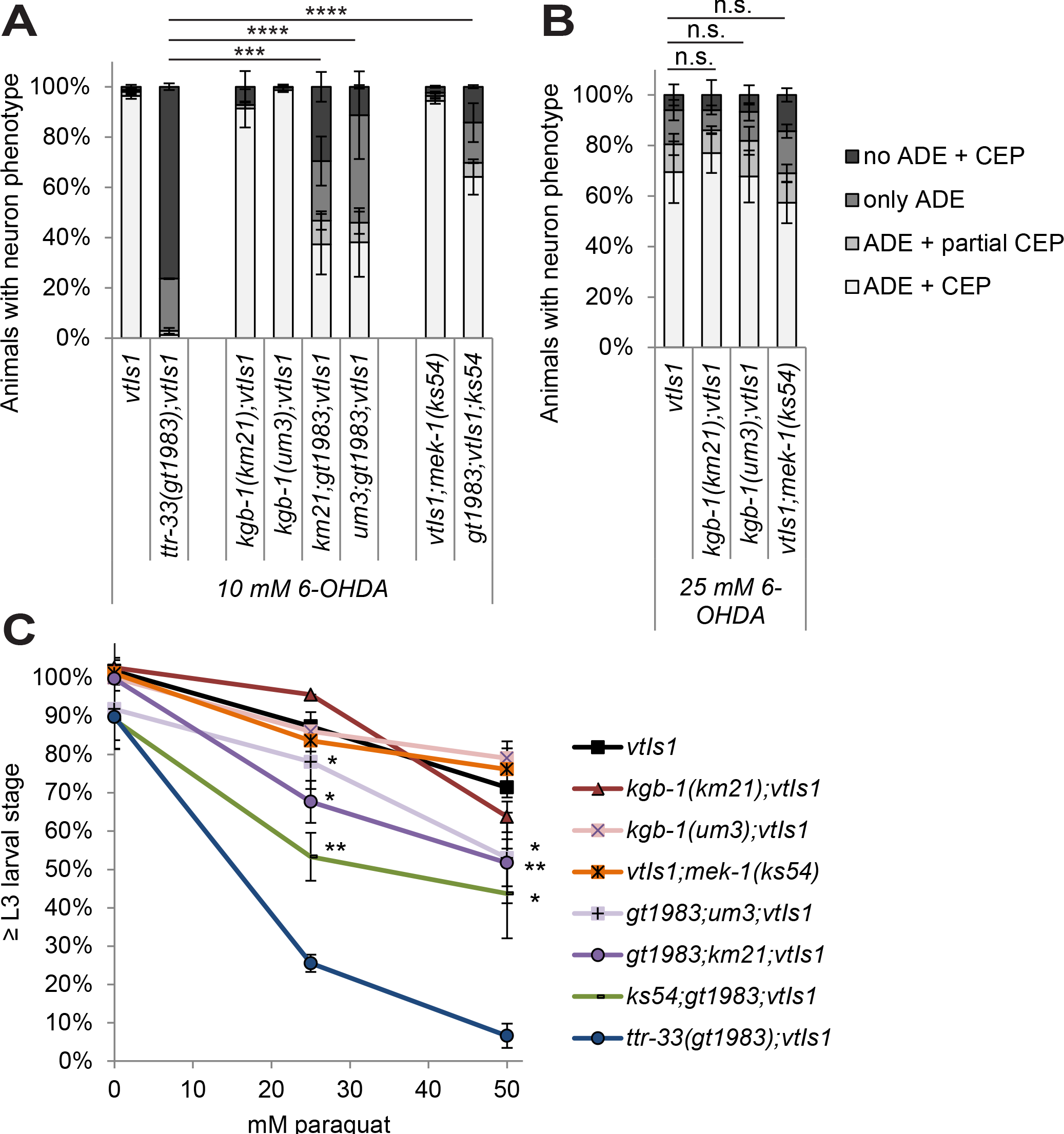
Interference with the *kgb*-*1* MAPK pathway decreases oxidative stress sensitivity in *ttr*-*33* mutants. (A) Effect of *kgb*-*1* MAP kinase pathway mutations on dopaminergic neurodegeneration after treatment with 10 mM 6-OHDA. Error bars = SEM of 3 biological replicates, each with 90-115 animals per strain. Total number of animals per strain n = 300-340 (****p<0.0001, ***p<0.001; G-Test). (B) Effect of *kgb*-*1* and *mek*-*1* MAP kinase pathway mutation on dopaminergic neurodegeneration after treatment with 25 mM 6-OHDA. Error bars = SEM of 3 biological replicates, each with 50-130 animals per strain. Total number of animals per strain n = 255-330 (n.s. p>0.05; G-Test). (C) Percentage of larvae reaching the L3 larval stage 24 hours after an 1 hour incubation with indicated concentration of paraquat. Error bars = SEM of 3-4 biological replicates for 25 and 50 mM paraquat and 2 biological replicates for 0 mM paraquat, each with 70-270 animals per strain and concentration. Total number of animals per condition n = 220-650 (**p<0.01, *p<0.05; two-tailed t-test comparing *ttr*-*33(gt1983)* double mutants to *ttr*-*33(gt1983)* single mutant data).

## Discussion

We report that the *C. elegans* transthyretin-related protein TTR-33 protects against 6-OHDA-induced dopaminergic neurodegeneration as well as against oxidative stress at an organismal level. The KGB-1 MAP kinase pathway mediates increased sensitivity to oxidative stress in the *ttr*-*33* mutant, whereas the JNK-1 MAP kinase pathway provides additional protection. In addition, we find that the corpse engulfment pathway is required for 6-OHDA-mediated dopaminergic neurodegeneration.

We speculate that TTR-33 is part of an endocrine defence system against oxidative stress. Transthyretin-related proteins form a nematode-specific expanded protein family that comprises 59 members, 54 of which are predicted to be secreted [24]. Some transthyretin-related genes are induced in response to environmental challenges such as oxidative stress or pathogen exposure: Oxidative stress induced by tert-butyl hydrogen leads to upregulation of several transthyretin-related genes [35], and treatment with the mitochondrial complex I inhibitor rotenone leads to the upregulation of 42 of the 59 transthyretin-related genes [36]. Furthermore, bacterial infection was reported to cause upregulation of *ttr*-*33* [37] and other transthyretin-related proteins [38]. However, we found that TTR-33 does not appear to be induced in response to oxidative stress and thus might provide constitutive protection. TTR-33 possesses 4 cysteine residues that are conserved within the *C. elegans* transthyretin-related protein family (marked in yellow in S2B Fig and S11 Fig). These cysteines could serve as reactive oxygen sensors or scavengers, analogous to those in the antioxidant glutathione. TTR-33 is likely produced in the six posterior arcade cells, which form a syncytium and are located between outer epidermis and the pharynx [32] and TTR-33 could be secreted from the arcade cells into the pharynx. This hypothesis is consistent with the partial rescue we observed upon expressing TTR-33 from an arcade cell-specific promoter. In the pharynx, TTR-33 could provide organismal protection against ingested oxidative stress-producing compounds. It will be interesting to experimentally investigate the mechanistic details of TTR-33 function in the future. *ttr*-*33* gene expression correlates with genes that are associated with the gene ontology terms ‘defence response to fungus’, ‘negative regulation of metalloenzyme activity’ and ‘neuropeptide signalling pathway’ (S1 Table). *ttr*-*33* mutants die prematurely and are hypersensitive to oxidative stress at an organismal level, supporting the idea that these animals might exhibit a compromised organismal defence system (reviewed in [39]). Alternatively, mutation of *ttr*-*33* could generally lower the threshold of oxidative damage both for dopaminergic neurons as well as at an organismal level.

We hypothesise that the KGB-1 and the JNK-1 MAPK pathways influence 6-OHDA-mediated neurodegeneration in the *ttr*-*33* mutant based on their role in organismal stress responses. KGB-1 is ubiquitously expressed [40,41] and was reported to transmit a neuroendocrine signal that mediates a food aversion behaviour upon disruption of core cellular processes such as translation, respiration and protein turnover [42]. It is conceivable that KGB-1 acts upstream of TTR-33. However, the *kgb*-*1* mutant on its own does not show enhanced 6-OHDA-induced neuronal loss. It is thus possible that the KGB-1 pathway senses the pathology conferred by the combination of exposure to oxidative stress and TTR-33 deficiency analogous to its described roles in sensing cellular defects [42]. Such sensing mechanisms might promote the degeneration of damaged neurons. In contrast, the JNK orthologues JNK-1 protects against 6-OHDA-induced neurodegeneration in the *ttr*-*33* mutant. This phenotype is consistent with the reported role of JNK-1 in protecting against oxidative stress [43,44].

The *C. elegans* corpse engulfment pathway could promote 6-OHDA-induced dopaminergic neurodegeneration by driving the phagocytosis of dopaminergic neurons. Morphological evidence for 6-OHDA-induced engulfment of dopaminergic neurons were reported [13,45]. However, it is not known exactly how dopaminergic neurons die. The engulfment pathway typically acts by eliminating apoptotic cell corpses, but has also been shown to promote cell death; mutations in the engulfment pathway enhance *C. elegans* apoptosis defects [15,17] and a small number of cell deaths are mediated by cell killing conferred by the engulfment machinery [18,19,46]. Consistent with our findings, reduced axonal degeneration in neurons whose axons have been severed has been observed in mutants with a defective engulfment machinery [47]. It is still poorly understood how dopaminergic neurons die in Parkinson’s disease patients. Active phagocytic cells in the brain are observed in almost all neurological diseases including Parkinson’s disease (for review [48,49]), as well as in 6-OHDA models of Parkinson’s disease (for review [50]). The *C. elegans* 6-OHDA model, in which the conserved engulfment pathway has a role in cell removal, might recapitulate important features of this disease.

## Methods

### *C. elegans* strains and maintenance

Strains were grown at 20°C unless indicated otherwise. Additional information about alleles in this study is provided by WormBase (www.wormbase.org) [51]. Strains from the *Caenorhabditis* Genetics Center (CGC) or the National BioResource Project (NBRP) were outcrossed to the BY200 strain to eliminate unlinked mutations and to introduce a green fluorescent protein (GFP) marker for dopaminergic neurons. The strain carrying the second *ttr*-*33* allele *gk567379* (generated as part of the Million Mutation project [52]) was identified with WormBase [51] and ordered from CGC. Both The isolated *gt1983* mutant and the *ttr*-*33(gk567379)* mutant were outcrossed a minimum of six times to the BY200 strain.

**BC15279** *dpy*-*5(e907)I; sEx15279*. sEx15279 [rCes F40C5.3::GFP + pCeh361] [53]
**BY200** *vtIs1[pdat-1::gfp; rol-6]* V (while *pdat-1::gfp* is always expressed, the penetrance of the roller phenotype is very low, only rarely detectable)
**CZ10175** *zdIs5 [Pmec-4::GFP]*
**N2 Bristol**
**QH6342** *zdIs5* I; *ttr*-*33(gt1983)* V
**TG2395** *cat*-**2(e1112)** II; *vtIs1* V;
**TG2400** *dat*-*1(ok157)* III; *vtIs1* V
**TG4103** *ttr*-*33(gt1983)* V; *vtIs1* V
**TG4104** *ttr-33(gk567379)* V; *vtIs1* V
**TG4106** *dat-1(ok157)* III; *ttr-33(gt1983)* V; *vtIs1* V
**TG4120** *pmk*-*1(km25)* IV; *vtIs1* V
**TG4121** *pmk*-*1(km25)* IV; *ttr*-*33(gt1983)* V; *vtIs1* V
**TG4122** *jnk*-*1(gk7)* IV; *vtIs1* V
**TG4123** *jnk*-*1(gk7)* IV; *ttr-33(gt1983)* V; *vtIs1* V
**TG4124** *pmk*-*3(ok169)* IV; *vtIs1* V
**TG4125** *pmk*-*3(ok169)* IV; *ttr-33(gt1983)* V; *vtIs1* V
**TG4126** *kgb*-*1(km21)* IV; *vtIs1* V;
**TG4154** *ced*-*2(e1752)* IV; *vtIs1* V;
**TG4155** *ced*-*2(e1752)* IV; *ttr-33(gt1983)* V; *vtIs1* V;
**TG4156** *ced*-*6(n1813)* III; *vtIs1* V;
**TG4157** *ced*-*6(n1813)* III; *ttr-33(gt1983)* V; *vtIs1* V;
**TG4158** *ced*-*10(n3246)* IV; *vtIs1* V;
**TG4159** *ced*-*10(n3246)* IV; *ttr-33(gt1983)* V; *vtIs1* V;
**TG4160** *ttr*-*52(sm211)* III; *vtIs1* V;
**TG4161** *ttr*-*52(sm211)* III; *ttr-33(gt1983)* V; *vtIs1* V;
**TG4170** *unc*-*119(ed3)* III; *cxTi10816* IV; Ex[Pttr-*33::gfp::3’ttr*-*33*, *Pmyo*-*2::mCherry*, *Pmyo*-*3::mCherry*]
**a**
**TG4171** vtIs1 V; CB4856
**TG4281** *unc*-*119(ed3)* III; *cxTi10816* IV; Ex[Pttr-*33::gfp::3’ttr*-*33*, *Pmyo*-*2::mCherry*, *Pmyo*-*3::mCherry*]
**b**
**TG4282** *unc*-*119(ed3)* III; *cxTi10816* IV; Ex[Pttr-*33::gfp::3’ttr*-*33*, *Pmyo*-*2::mCherry*, *Pmyo*-*3::mCherry*]
**c**
**TG4283** *kgb*-*1(km21)* IV; *ttr*-*33(gt1983)* V; vtIs1 V
**TG4284** *kgb*-*1(um3)* IV; *ttr*-*33(gt1983)* V; vtIs1 V
**TG4285** *ttr*-*33(gt1983) V;* vtIs1 V; *mek*-*1(ks54)* X
**TG4286** *kgb*-*1(um3)* IV; *vtIs1* V;
**TG4336** *ttTi5605* II; *Is[Pbath*-*15::ttr*-*33::3’ttr*-*33*, *unc*-*119(+)*, integration into *ttTi5605* MosSCI site/ II; *unc*-*119(ed9)* III
**TG4337** *ttTi5605* II; *Is[Pttr*-*33::ttr*-*33::3’ttr*-*33*, *unc*-*119(+)*, integration into *ttTi5605* MosSCI site] II; *unc*-*119(ed9)* III
**TG4338** *ttTi5605* II; *unc*-*119(ed9)* III. *Ex[Pbath*-*15::ttr*-*33::3’ttr*-*33*, *unc*-*119(+)*, *Pmyo*-*2::mCherry*, *Pmyo*-*3::mCherry] a*
**TG4339** *ttTi5605* II; *unc*-*119(ed9)* III. *Ex[Pbath*-*15::ttr*-*33::3’ttr*-*33*, *unc*-*119(+)*, *Pmyo*-*2::mCherry*, *Pmyo-3::mCherry] b*
**TG4340** *ttTi5605* II; *unc*-*119(ed9)* III. *Ex[Pbath*-*15::ttr*-*33::3’ttr*-*33*, *unc*-*119(+)*,*Pmyo*-*2::mCherry*, *Pmyo*-*3::mCherry] c*
**TG4341** *ttTi5605* II; *unc-119(ed9)* III. *Ex[Pttr*-*33::ttr*-*33::3’ttr*-*33*, *unc*-*119(+)*, *Pmyo*-*2::mCherry*, *Pmyo*-*3::mCherry] a*
**TG4342** *ttTi5605* II; *unc*-*119(ed9)* III. *Ex[Pttr*-*33::ttr-33::3’ttr*-*33*, *unc*-*119(+)*, *Pmyo*-*2::mCherry*, *Pmyo*-*3::mCherry] b*
**TG4343** *ttTi5605* II; *unc*-*119(ed9)* III. *Ex[Pttr*-*33::ttr*-*33::3’ttr*-*33*, *unc*-*119(+)*, *Pmyo*-*2::mCherry*, *Pmyo*-*3::mCherry] c*

### Mutagenesis and mapping

L4 and young adult staged BY200 animals were mutagenised with 25 mM ethyl methanesulfonate in M9 buffer for 4 hours at 20°C, washed and grown at 15°C overnight on NGM plates seeded with OP50 *E. coli*. The animals were bleached once they started to lay eggs and the F1 progeny left to hatch overnight in M9. This synchronised F1 population was left to develop on seeded plates until reaching early adult stage. The F1 adults were then washed off in M9 to lay a synchronised population of F2 L1 larvae which were then treated with 10 mM 6-OHDA and screened for dopaminergic neuron loss after 72 hours. Candidates were picked on separate plates, backcrossed at least three times and re-tested for their hypersensitivity to 6-OHDA. The genomic DNA was sent for whole-genome sequencing and SNP mapping was performed as previously described [28,29]. SNP mapping combined with whole-genome sequencing revealed that the location of the phenotype-causing mutation could be narrowed down to an interval between 4 and 9 Mb on chromosome V. This region, besides *ttr*-*33*, contained two other exon-altering mutations that were excluded by testing a deletion allele (*fbxa*-*223*) and by fosmid injection *(F43H9.4). ttr*-*33* identity was further confirmed by genetic rescue and by complementation testing using the second allele *ttr*-*33(gk567379)* which was obtained from the CGC.

### *C. elegans* assays

All assays were performed in a blinded manner.

For **6-OHDA assays**, 1-10 adult animals were incubated to lay eggs in 70μL M9 without food on a shaker at 20°C, 500 rpm for 24-30 h. 10 μL 200 mM ascorbic acid and 10 μL of the respective 6-OHDA 5x stock concentration were freshly prepared in H_2_O and added to 30 μL of L1-stage larvae in M9. 6-OHDA concentrations were chosen such that differential sensitives of various strains could be efficiently analysed. After 1 hour incubation at 20°C and shaking at 500 rpm, 150 μL M9 buffer was added to oxidise and inactivate the 6-OHDA. The animals were pipetted to one half of an NGM plate containing a stripe of OP50 bacteria on the opposite half of the plate and adult animals and eggs were picked off to prevent growth of those that were not treated at L1 stage. Plates were incubated at 20°C before examining the 6-OHDA-induced degeneration using a Leica fluorescent dissecting microscope. Unless otherwise indicated, head dopaminergic neurons (ADE and CEP neurons) were scored 72 hours after treatment in technical duplicates. <u>P</u>osterior <u>d</u>eirid (PDE) dopaminergic neurons, which are formed later in L2 stage larvae, did not appear to be affected by 6-OHDA treatment, as previously observed [13]. For treatment of different developmental stages, the respective stages were picked after 6-OHDA treatment from a mixed population of animals.

For **paraquat and hydrogen peroxide (H_2_O_2_) assays**, 10 μL of the 5x paraquat or H_2_O_2_ stock concentration was freshly prepared and added to 40 μL of filtered L1-stage larvae in M9. After 1 hour incubation in the shaker, 150 μL M9 buffer was added and the L1 larvae were counted after pipetting approximately 100 of them to a seeded NGM plate. After 24 hours, animals that reached L3 larval stage were examined and scored.

For **laser axotomy and microscopy**, **a**nimals were immobilised in 0.05% tetramisole hydrochloride on 3.5% agar pads. Axotomies were performed as previously described [54], using a MicroPoint Laser System Basic Unit attached to a Zeiss Axio Imager A1 (Objective EC Plan-Neofluar 100X/1.30 Oil M27). Axons were severed in L4 stage animals 50 μm anterior to the PLM cell body, visualised on a Zeiss Axio Imager Z1 microscope equipped with a Photometrics camera (Cool Snap HQ2; Tucson, AZ) and analysed with MetaMorph software (Molecular Devices, Sunnyvale, CA) or Image J. Animals were analysed 24 hours post-axotomy for regrowth, reconnection and fusion. Regrowth was recorded as the length of regenerative growth from the proximal side of the cut site: the proximal axon was traced from the start of regrowth to the tip of the longest regenerative branch using Metamorph software; neurons that underwent axonal fusion were excluded from these quantifications. Axons were deemed to be reconnected when the proximal and distal axons appeared connected and within the same focal plane when observed with a 63X objective. Axonal fusion was defined as proximal-distal reconnection that prevented the onset of degeneration in the distal axon. For clarification of successful fusion, animals were re-analysed at 48 hours and/or 72 hours post-axotomy as necessary.

For **lifespan assays**, 50 L4 stage animals were picked for each strain and moved to new plates every day during the first 6 days and again at later days to avoid mixing with the progeny. Animals that did not move after being touched with a platinum pick were scored as dead. Dead animals were counted and removed every day. ‘Bag of worms’ and dry and burst animals were not included in the lifespan analysis. The data was analysed using OASIS (Online application for Survival Analysis, http://sbi.postech.ac.kr/oasis/) [55].

For **egg-laying and development assays**, over 20 L4 stage animals per strain were picked separately 24 hours before the assay. On the day of the assay, two plates per strain were seeded with concentrated OP50 *E. coli* and ten adults each deposited onto the plates. The adults were left to lay eggs for 1.5 hours, removed from the plate and the eggs counted. Developmental stages were determined at 48 and 61 hours, respectively, after egg-laying.

**Dopamine paralysis assays** were carried out as previously described [56]. Well-fed adults 24 hours post the L4 stage were incubated for 20 min on a plate containing different concentrations of dopamine. Animals were scored as paralysed when they failed to complete a full sinusoidal body movement during a 5 second observation period. Two plates containing 25 animals each were assessed for each condition.

**Basal slowing assays** were performed as previously described [57]. NGM plates were prepared and half of them seeded with HB101 *E. coli* before incubating them at 37°C overnight. Also, mid-L4 larvae were picked separately on seeded plates. The next day, these staged, well-fed young adult animals were placed in 40 μL of M9 for 2 minutes to clean them from bacteria, and then picked to the centre of a plate. 6 animals each were put on a plate with and without bacteria. After 2 minutes adaptation time, the locomotion rate of each animal was quantified by counting the number of body bends completed in five consecutive 20 s intervals. The experiment was performed twice on different days.

### qRT-PCR

L1 stage animals were filtered using an 11 μm filter from two 9 cm plates containing mixed stage animals. Animals were washed twice and the population was split in half for control treatment and drug treatment. For the 6-OHDA assays, ascorbic acid and H_2_O was added to the control wells and 10 mM 6-OHDA to the treated animals. For paraquat assays, H_2_O was added to the control wells and 25 mM paraquat to the treated animals. After 1 hour of treatment, RNA extraction was performed based on [58] using Trizol (Thermo Fisher Scientific). The following changes were introduced. The chloroform extraction was repeated once more and 0.1 volumes of 3 M sodium acetate was added to the propanol extraction to help RNA precipitation. The RNA pellet was resuspended in 15 μL H_2_O. The gDNA was digested using the Turbo DNA-free kit (Ambion Life Technologies) in a total volume of 18 μL. 1 μg of RNA was reverse transcribed to cDNA with 15 bp oligo dT primers using the M-MLV Reverse Transcriptase (Promega) according to manufacturers’ instructions. The cDNA was analysed by qRT-PCR using the StepOnePlus Real-Time PCR System (Applied Biosystems) applying melting temperature of 62°C. The following primers were designed using QuantPrime [59]: *ttr*-*33*/C37C3.13.2_F: ACCCAATCAGCTGGAGTTAAGGG and ttr-33/C37C3.13.2_R: ATCCGGTCCGGTATCATCATCG, *gst*-*4*/K08F4.7_F: ATGGTCAAAGCTGAAGCCAACG and *gst*-*4*/K08F4.7_R: ACTGACCGAATTGTTCTCCATCG, *gcs*-*1*/F37B12.2.1_F: GTTGATGTGGATACTCGGTGTACG and *gcs*-*1*/F37B12.2.1_R: ATCTCTCCAGTTGCTCGTTTCG, *gst*-*1*/R107.7.1_F: CGTCATCTCGCTCGTCTTAATGGG and *gst*-*1*/R107.7.1_R: TGGTGTGCAAATCACGAAGTCC, Y45F10D.4_F: TTGCCT CAT CTT CCCTGGCAAC and Y45F10D.4_R: GT CTT GGGCGAGCATT GAACAG, *pmp*-*3*/C54G10.3b_F: ACTTGCTGGAGTCACTCATCGTG and *pmp*-*3*/C54G10.3b: TCAAGGCAATCCGAGTGAAGACG. The PCR mixture consisted of 0.2 μM primers, cDNA (diluted 1:50) and 1× Power SYBR Green PCR Master Mix (Thermo Fisher Scientific). All reactions were run in technical triplicates. Relative abundance was calculated using the ΔΔCt method [60]. To control for template levels, values were normalised against the control genes *Y45F10D.4* and *pmp*-*3* [61,62].

### Bioinformatics analysis

The **open reading frame and exon-intron structure** of *ttr*-*33* was derived from information compiled in WormBase (www.wormbase.org) [51]. **Extra- and intracellular and transmembrane protein domains and signal peptides** were predicted with Phobius (http://phobius.sbc.su.se/) [63]. The **protein structure model** was calculated in SWISS-MODEL (swissmodel.expasy.org) [64] and visualised with Swiss-PdbViewer 4.1.0 (http://www.expasy.org/spdbv/) [65]. Jalview Version 2.9.0b2 (www.jalview.org) was used for phylogenetic alignment (ClustalWS version 2.0.12 with default settings) [66]. Secondary structure prediction was performed with JPred4 [67] in Jalview [68]. For the correlated gene expression analysis, 100 result genes calculated by SPELL (Serial Pattern of Expression Levels Locator) Version 2.0.3.r71 (http://spell.caltech.edu:3000/) were considered. Given a query gene, SPELL searches for genes with correlated expression patterns based on available microarray, RNAseq and tiling array data (currently 5264 experiments in 315 datasets). The main algorithm is described in [69].

### Statistical analysis

The two-tailed t-test was performed with the Analysis ToolPak Add-In in Microsoft Excel 2010. To choose between a t-test assuming equal or unequal variances, an F-test was conducted with the Analysis ToolPak Add-In. A **G-Test** was performed with R Studio 1.0.44 DescTools descriptive statistics tools package to compare 6-OHDA treatment data. p-values were calculated for each biological replicate and for the pooled biological replicate data, and the least significant of these p-values is indicated in the graphs. Data is only marked as statistically significant if all replicates and the pooled replicate data were found to be significant. Statistical analysis for axonal regeneration was performed using GraphPad Prism and Microsoft Excel. Two-way comparison was performed using the *t*-test.

### DNA cloning and generation of transgenic lines

For generation of transgenic constructs, genomic DNA was amplified using primers containing additional 8 base pair restriction sites (marked in italics) for insertion into the pCFJ151 and the pCFJ212 vector for insertions on chromosome II and IV, respectively [70]. In the following primer sequences, restriction sites are highlighted in italics, translation start and translation stop codons in bold. Primers for the transcriptional reporter construct *Ascl* – Pttr-33 – *NotI* – GFP – *Fsel* – 3′ttr-33 – *Pacl*: AscI_Pttr-33 – GTACTT*ggcgcgcc*CAAAGAAACTTGCGTGTTTC, Pttr-33_Notl – AAGGAAAAAAgcggccgca**CAT**TATTTTT GTCT GAAAAT ACAAACAC, Notl_GFP – GCTA*gcggccgc*AGTAAAGGAGAAGAACTTTTCACTGG, GFP_FseI – TTATTAggccggccCCTTGTATGGCCGGCTAG, Fsel_3'ttr-33 – ACAAGG*ggccggcc***TAA**TAATTATACTTAAAAGTATTTTCAACTG, 3'ttr-33_Pacl – GCTA*ttaattaa*GCATGCATCTTCCCATAAAAG. Additional primers for the translational construct *Ascl* – Pttr-33 – *NotI* – TTR-33 – *FseI* – 3'ttr-33 – *Pacl*: Notl_TTR-33 – AAGGAAAAAA*gcggccgc*TCACGACTCGCGTGCATT, TTR-33_FseI – AGAAAA*ggccggcc*tTCTGTGAAGACAGTCACGTGACTC. The *Pttr*-*33* promoter sequence in these constructs ranges from -1500 to the translational start site. The predicted promoter sequence of *ttr*-*33* ranges from -499 to 100 bp with respect to the translational start site [71]. Additional primers for the translational construct *AscI* – Pbath-15 – *NotI* – TTR-33 – *FseI* – 3′bath-15 – *PacI:* AscI_Pbath-15: GTACTT*ggcgcgcc*TTTTCACTTTTTTTGGTCAGAATC, Pbath-15_Notl: AAGGAAAAAA*gcggccgcc***CAT**IIIGAAGAGTTTGCTG, FseI_3′bath-15: CTCAT*Aggccggcc***TAA**TCATTTTTTTCCTGTTTTAGC, 3'bath-15_Pacl: GAGACC*ttaattaa*CTGGAGTTCACCACTACAAC. For single-copy integration of transgene according to [70], the constructs were injected into *unc-119* animals, together with one or two fluorescent marker(s) (*Pmyo-2::mCherry* to label the pharynx and *Pmyo-3::mCherry* to label the bodywall), and the Mos transposase under the control of the *eft*-*3* promoter as described [72]. Wild-type moving animals that did not show expression of fluorescent markers were singled out and tested for transgene integration by PCR. At least three independent transgenic lines were analysed for all translational and transcriptional constructs.

## Acknowledgements

We thank Neda Masoudi for advice at initial stages of the project, Radoslav Lukoszek for advising on qRT-PCR experiments and Ulrike Gartner for proofreading. Some strains were provided by the CGC, which is funded by NIH Office of Research Infrastructure Programs (P40 OD010440).

## Supporting information

### S1 Fig. Effect of single-copy transgenes on 6-OHDA hypersensitivity of *ttr*-*33* mutant

Effect of *Pttr-33::ttr*-*33* and *Pbath-15::ttr*-*33* single copy integration on dopaminergic neurodegeneration of *ttr*-*33* mutant animals after treatment with 10 mM 6-OHDA. Error bars = SEM of 2 biological replicates, each with 50-105 animals per strain and concentration. Total number of animals per condition n = 150-205 (n.s. p>0.05; G-Test).

### S2 Fig. TTR-33 protein structure prediction

(A) TTR-33 signal peptide (in red) and non-cytoplasmic part (in blue) as determined using the Phobius webserver [63] (http://phobius.sbc.su.se/). No transmembrane domains (in grey) and no cytoplasmic parts (in green) were predicted. (B) TTR-33 secondary structure with helices (red tubes) and beta strands (green arrows) as predicted by JPred4 [67]. (C) Predicted TTR-33 structure (in grey with indicated mutated sites in red and cysteine bridges in yellow) with superimposed TTR-52 structure (in black). The small TTR-52 α-helix (top right part of structure) is likely an artefact caused by mutations that were introduced for crystallisation [25].

### S3 Fig. Development and dopamine-mediated behaviours of *ttr*-*33* mutants

(A) Developmental stages of wild-type and *ttr*-*33* mutant embryos 48 and 61 hours after egg-laying. Error bars = SEM of 3 biological replicates, each with 40-230 animals per strain. Total number of animals n = 405-515 (*p<0.05, n.s. p>0.05; G-Test). (B) Dopamine paralysis assay: Ability of young adult animals to move on plates with indicated concentrations of dopamine. Error bars = StDev of 2 technical replicates, each with 25 animals per strain and condition. Total number of animals per condition n = 50. (n.s. p>0.05; two-tailed t-test comparing wild-type and mutant animal data at 100 mM dopamine). (C) Basal slowing response: Ability of young adult animals to slow down on a lawn of bacteria. The tyrosine hydroxylase mutant *cat*-*2 (abnormal <u>cat</u>echolamine distribution)* is deficient of dopamine synthesis. The values for each animal are depicted with grey symbols and the average across animals is indicated with a black bar. Error bars = SEM of 2 biological replicates, each with 6 animals per strain and state. Total number of animals per condition n = 12 (****p<0.000001, *p<0.05; two-tailed t-test).

### S4 Fig. 6-OHDA concentration gradient and effect of *ttr-52* mutations on dopaminergic neurodegeneration

(A) Dopaminergic neurodegeneration in wild-type and *ttr*-*33(gt1983)* mutant animals after treatment with a distinct concentrations of 6-OHDA. Error bars = SEM of 2 biological replicates, each with 75-120 animals per strain. Total number of animals per condition n = 180-220. (B) Effect of *ttr*-*52* mutation on dopaminergic neurodegeneration after treatment with 10 mM 6-OHDA. Error bars = SEM of 2 biological replicates, each with 65-110 animals per strain. Total number of animals per condition n = 165-220 (n.s. p>0.05; G-Test). (C) Effect of *ttr*-*52* mutation on dopaminergic neurodegeneration after treatment with 50 mM 6-OHDA. Error bars = SEM of 4 biological replicates, each with 100-125 animals per strain. Total number of animals per condition n = 430-440 (n.s. p>0.05; G-Test).

### S5 Fig. Effect of mutations in the cell corpse engulfment pathway on dopaminergic neurodegeneration

(A) Effect of mutations in engulfment pathway genes on dopaminergic neurodegeneration after treatment with 2.5 mM 6-OHDA. Error bars = SEM of 2 biological replicates, each with 100-110 animals per strain. Total number of animals per condition n = 200-220 (****p<0.0001, n.s. p>0.05; G-Test). (B) Effect of mutations in engulfment pathway genes on dopaminergic neurodegeneration after treatment with 25 mM 6-OHDA. Error bars = SEM of 2 biological replicates, each with 85-140 animals per strain. Total number of animals per condition n = 205-230 (****p<0.0001, ***p<0.001, **p<0.01, n.s. p>0.05; G-Test).

### S6 Fig. *ttr*-*33* mutant lifespan

Lifespan data for second biological replicate including 85-110 animals per strain. The inset shows the mean lifespan with the error bars depicting the standard error (****Bonferroni p≤0.0001; Log-Rank Test).

### S7 Fig. Transcript levels of *ttr*-*33*, *gst*-*4*, *gcs*-*1* and *gst*-*1* normalised against the control gene *Y45F10D.4*

(A) *ttr-33,* (B) *gst-4,* (C) *gcs-1* and (D) *gst*-*1* mRNA levels in wild-type and *ttr*-*33* mutant L1 stage larvae after treatment with 10 mM 6-OHDA. (E) *ttr*-*33*, (F) *gst*-*4*, (G) *gcs*-*1* and (H) *gst*-*1* mRNA levels in *ttr*-*33* mutants at L1 stage as compared to wild-type larvae under control conditions (no 6-OHDA exposure). (I) *ttr*-*33*, *gst*-*4*, *gcn*-*1* and *gst*-*1* mRNA levels in wild-type and *ttr*-*33* mutant L1 stage larvae after treatment with 25 mM paraquat. (A)-(I) The data are normalised to the control gene *Y45F10D.4* [61,62]. The average and the respective values for the biological replicates (biorep a-f) are indicated. Error bars = SEM of 5-6 biological replicates (n.s. p>0.05).

### S8 Fig. Transcript levels of *ttr*-*33*, *gst*-*4*, *gcs*-*1* and *gst*-*1* normalised against the control gene *pmp*-*3*

(A) *ttr*-*33*, (B) *gst*-*4*, (C) *gcs*-*1* and (D) *gst*-*1* mRNA levels in wild-type and *ttr*-*33* mutant L1 stage larvae after treatment with 10 mM 6-OHDA. (E) *ttr*-*33*, (F) *gst*-*4*, (G) *gcs*-*1* and (H) *gst*-*1* mRNA levels in *ttr*-*33* mutant at L1 stage as compared to wild-type larvae under control conditions (no 6-OHDA exposure). (I) *ttr*-*33*, *gst*-*4*, *gcn*-*1* and *gst*-*1* mRNA levels in wild-type and *ttr*-*33* mutant L1 stage larvae after treatment with 25 mM paraquat. (A)-(I) The data are normalised to the control gene *pmp-3* (*C54G10.3b*) [61,62]. The average and the respective values for 5-6 biological replicates (biorep a-f) are indicated. Error bars = SEM of 5-6 biological replicates (**p<0.01, *p<0.05; n.s. p>0.05; two-tailed t-test).

### S9 Fig. Effect of MAPK pathway mutations on dopaminergic neurodegeneration

(A) Effect of p38 and JNK stress response pathway mutations on dopaminergic neurodegeneration after treatment with 10 mM 6-OHDA. Error bars = SEM of 2-3 biological replicates, each with 100-115 animals per strain and concentration. Total number of animals per condition n = 100-315 (***p<0.001, n.s. p>0.05; G-Test). (B) Effect of p38 and JNK stress response pathway mutations on dopaminergic neurodegeneration after treatment with 50 mM 6-OHDA. Error bars = SEM of 3 biological replicates, each with 100-110 animals per strain. Total number of animals per strain n = 305-320 (n.s. p>0.05; G-Test).

### S10 Fig. Effect of *kgb*-*1* MAPK pathway mutations on dopaminergic neurodegeneration and development

(A) Effect of *kgb*-*1* MAP kinase pathway mutations on dopaminergic neurodegeneration after treatment with 2.5 mM 6-OHDA. Error bars = SEM of 2-3 biological replicates, each with 95-120 animals per strain. Total number of animals per strain n = 200-340 (n.s. p>0.05; G-Test). (B) Effect of *kgb*-*1* MAP kinase pathway mutations on dopaminergic neurodegeneration after treatment with 50 mM 6-OHDA. Error bars = SEM of 3 biological replicates, each with 100-125 animals per strain. Total number of animals per strain n = 205-335 (n.s. p>0.05; G-Test). (C) Number of laid eggs per animal per hour. Error bars = SEM of 3-4 biological replicates, each with 20 animals per strain. Total number of animals n = 60-80 (n.s. p>0.05; two-tailed t-test). (D) Developmental stages of wild-type and mutant embryos 48 hours after egg-laying. Error bars = SEM of 3 biological replicates, each with 40-125 animals per strain. Total number of animals n = 150-350 (**p<0.01; two-tailed t-test).

### S11 Fig. Alignment of *C. elegans* transthyretin-related proteins

Amino acids above an identity threshold of 69 are coloured according to their physicochemical properties using the Zappo colour code. Conservation score, quality and consensus are each shown as a bar graph in the bottom of the alignment.

**S1 Table.** Gene Ontology (GO) term analysis for genes for which the expression pattern correlated with *ttr*-*33*. Genes with expression patterns correlating to the *ttr*-*33* expression pattern were predicted using SPELL [69] – The GO terms and the according genes are sorted by p-value. The number of genes that are associated with this GO term out of all genes with a similar expression pattern to *ttr*-*33*, and out of all genes in the genome, are shown in column ‘% Query’ and ‘% Genome’, respectively. Very specific processes that are associated with few genes in the genome are marked in blue. * Similar GO terms based on the same group of genes were merged.

| GO Term | p-value | % Query | % Genome | Annotated genes |
| --- | --- | --- | --- | --- |
| neuropeptide signaling pathway | 5.21E-08 | 7 of 41 | 188 of 54514 | NLP-31, FLP-9, FLP-21, FLP-24, NLP-15, NLP-27, FLP-22 |
| (negative) regulation of metalloenzyme activity, (negative) regulation of membrane protein ectodomain proteolysis* | 2.73E-04 | 2 of 41 | 2 of 54514 | CRI-2, K07C11.3 |
| single-multicellular organism process* | 3.79E-04 | 16 of 41 | 5401 of 54514 | FLP-24, F20A1.1, NAS-36, GRD-10, FLP-12, BEST-24, LET-522, CAL-2, SNET-1, TNC-2, RPS-20, Y73F4A.2, STA-2, T26C5.4, PTR-8, F40F12.7 |
| (defense) response to fungus * | 9.53E-04 | 3 of 41 | 32 of 54514 | NLP-31, CNC-6, STA-2 |
| multi-organism process | 1.51E-02 | 7 of 41 | 1212 of 54514 | NLP-31, FLP-21, F20A1.1, FLP-12, CNC-6, STA-2, C18E3.5 |
| biological regulation | 1.96E-02 | 23 of 41 | 14131 of 54514 | NLP-31, FLP-9, FLP-21, FLP-24, W01B11.6, NLP-15, NLP-27, GRD-10, FLP-12, BEST-24, TTR-48, CRI-2, LET-522, H05L03.3, TTR-32, SNET-1, K07C11.3, RPS-20, STA-2, PTR-8, FLP-22, RIC-4, F40F12.7 |
| locomotion | 3.28E-02 | 8 of 41 | 1873 of 54514 | NAS-36, NLP-15, GRD-10, BEST-24, LET-522, TTR-32, SNET-1, Y4C6B.2 |

**S4 Movie. Pttr-33::GFP expression in *C. elegans* embryo.**

**S5 Movie. Pttr-33::GFP expression in L2 stage *C. elegans* larva.**

**S1 Movie. Pttr-33::GFP expression in L3 stage *C. elegans* larva.**

**S2 Movie. Pttr-33::GFP expression in L4 stage *C. elegans* larva.**

**S3 Movie. Pttr-33::GFP expression in young adult *C. elegans.***

